# Investigating the performance of foundation models on human 3’UTR sequences

**DOI:** 10.1101/2024.02.09.579631

**Authors:** Sergey Vilov, Matthias Heinig

## Abstract

Foundation models, such as DNABERT and Nucleotide Transformer have recently shaped a new direction in DNA research. Trained in an unsupervised manner on a vast quantity of genomic data, they can be used for a variety of downstream tasks, such as promoter prediction, DNA methylation prediction, gene network prediction or functional variant prioritization. However, these models are often trained and evaluated on entire genomes, neglecting genome partitioning into different functional regions. In our study, we investigate the efficacy of various unsupervised approaches, including genome-wide and 3’UTR-specific foundation models on human 3’UTR regions. Our evaluation includes downstream tasks specific for RNA biology, such as recognition of binding motifs of RNA binding proteins, detection of functional genetic variants, prediction of expression levels in massively parallel reporter assays, and estimation of mRNA half-life. Remarkably, models specifically trained on 3’UTR sequences demonstrate superior performance when compared to the established genome-wide foundation models in three out of four downstream tasks. Our results underscore the importance of considering genome partitioning into functional regions when training and evaluating foundation models. In addition, the proposed set of 3’UTR-specific tasks can be used for benchmarking future models.

## Introduction

Foundation models, adopted from the field of Natural Language processing, have recently catalyzed a transformative shift in DNA research. Trained in a self-supervised manner on vast amounts of genomic data, these models offer a comprehensive understanding of the sequence language and hold the potential to enable various downstream applications. Genomic foundation models have achieved encouraging results in promoter (Ji *et al*., 2021; Dalla-Torre *et al*., 2023; Zhou *et al*., 2023) and enhancer (Dalla-Torre *et al*., 2023) prediction, chromatin state analysis (Lee *et al*., 2022), transcription factor binding sites prediction (Ji *et al*., 2021; Dalla-Torre *et al*., 2023; Zhou *et al*., 2023), and functional variants prioritization (Ji *et al*., 2021; Dalla-Torre *et al*., 2023).

The DNABERT model (Ji *et al*., 2021) was one of the first adaptations of large language models to the human genome. It was based on the transformer BERT architecture (Devlin *et al*., 2019), which is particularly suitable for modeling long-range interactions due to its self-attention mechanism. DNABERT outperformed previous state-of-the-art models in various downstream tasks, including promoter prediction and splice site recognition. However, trained solely on the human genome, DNABERT was not able to utilize evolutionary information, which has long been leveraged by alignment-based conservation models, such as PhyloP (Pollard *et al*., 2010) or PhastCons (Siepel *et al*., 2005). These models are commonly used directly or as a part of more complex models for numerous applications, including prioritization of functional variants. This limitation was addressed in the subsequent DNABERT-2 model (Zhou *et al*., 2023) which was trained on 135 species, enabling utilization of evolutionary information in an alignment-free manner. In parallel to DNABERT-2, another collection of transformer-based foundation models, called Nucleotide Transformer (NT) (Dalla-Torre *et al*., 2023), was developed. This collection was initially represented by NT-500M-human, NT-500M-1000g, NT-2.5B-1000g, and NT-2.5B-multi models, where “human,” “1000g,” and “multi” correspond to the GRCh38/hg38 human reference genome, 3202 high-coverage human genomes from the 1000 Genome project, and genomes from 850 distinct species, respectively. Subsequently, introduction of various architectural tricks permitted developing more compact variations of these models with comparable efficiency (v2-50M, v2-100M, v2-250M, and v2-500M).

These foundational models showed promising performance when trained and evaluated across the entire human genome. The attention mechanism plays a crucial role within the transformer model, enabling the comprehension of genomic context within the perception field. However, the perception field of modern foundation models is still limited to 1,000-100,000 base pairs (bp), which hinders their capacity to gain a comprehensive understanding of a wider genomic landscape, including the positioning of sequences in relation to various functional genomic regions like coding regions, promoters, enhancers, 3’UTRs, and so on. Yet, the published studies on genomic foundation models fall short in providing a thorough evaluation of these models on individual regions of the genome.

Another significant aspect that has received limited attention is the capacity of language models to identify regulatory elements that could exhibit mobility within the genomic sequence, such as binding motifs of RNA binding proteins (RBP). While prior research has demonstrated the use of BERT-like models for identifying such motifs in yeast (Gankin *et al*., 2023; Karollus *et al*., 2023), the extent to which language models can accomplish this task in the context of human genome remains unclear.

In this study, we compare the performance of alignment-free foundational models and established alignment-based conservation models on 3’UTR sequences of the human genome through a range of downstream tasks, including recognition of RBP binding motifs, prioritization of functional variants, prediction of expression levels in massively parallel reporter assays (MPRA), and estimation of mRNA half-life. To investigate potential advantages of models trained and evaluated on specific regions compared to the entire genome, we train 6 additional transformer models, 5 of which solely on 3’UTR sequences from 241 mammalian genomes.

## Methods

### Models

In total, 9 alignment-free and 3 alignment-based models were evaluated on human 3’UTR sequences, including 5 alignment-free models specifically trained on 3’UTRs.

#### Genome-wide transformer models

*DNABERT*

The first BERT-like model for DNA analysis. We used the version of DNABERT with 6-mer encoding for sequences (DNABERT-6) as it demonstrated the best performance (Ji *et al*., 2021).

*DNABERT-2*

BERT-like model trained on whole genome data from 135 species.

*DNABERT2-ZOO*

We retrain the original DNABERT-2 model based on the whole Zoonomia dataset (Zoonomia Consortium, 2020), described in detail in the next section.

*NT-MS-v2-250M*

250M-parameter Multispecies NT model. Although the 2.5B Multispecies NT model outperformed the other NT models across the downstream tasks (Dalla-Torre *et al*., 2023), its training on the available GPU infrastructure would be challenging due to a substantial number of learning parameters. We therefore opted for the v2-250M version which previously demonstrated close performance (Dalla-Torre *et al*., 2023).

#### 3’UTR-specific transformer models

*DNABERT-3UTR, DNABERT2-3UTR*, and *NTv2-250M-3UTR*

We retrain the previously proposed DNABERT, DNABERT-2, and NT-MS-v2-250M models solely on 3’UTR sequences from the Zoonomia dataset (Zoonomia Consortium, 2020).

*StateSpace* and *StateSpace-SA*

We adopted the state space model previously used for language modeling on 3’UTR sequences in yeasts (Gankin *et al*., 2023; Karollus *et al*., 2023). This model permits introducing species-awareness by including the corresponding species label as additional input (Gankin *et al*., 2023; Karollus *et al*., 2023). We trained both species-agnostic (*StateSpace*) and species-aware (*StateSpace-SA*) state space models.

Language models training on the Zoonomia dataset was performed for about 2 epochs on 10 NVIDIA A100 80G GPUs in parallel using the AdamW optimizer (Loshchilov and Hutter, 2019). To accelerate the training procedure, we used the mixed precision training and gradient accumulation techniques. The training parameters (Table S1) were set as close as possible to the original model publication. All the models were trained using the masked language model loss. For each input sequence, 15% of the tokens were randomly chosen for masking. Of these tokens, 80% were replaced with a masking token, 10% were randomly mutated, and the remaining 10% remained unchanged. Since a large fraction of sequences in the Zoonomia dataset contained long N-chunks, we added an extra ‘NNNNNN’ token when training the DNABERT-2 and NTv2-250M architectures.

#### Alignment-based conservation models

*PhyloP-100way*

Conservation model based on whole genome alignment of 100 vertebrates (Pollard *et al*., 2010).

*PhyloP-241way*

Conservation model based on whole genome alignment of 241 mammalian genomes from the Zoonomia project (Zoonomia Consortium, 2020).

*CADD-1*.*7*

Combined Annotation-Dependent Depletion model v. 1.7 (Schubach *et al*., 2024). This is a logistic regression-based classifier that directly predicts deleteriousness or functionness of a genetic variant in the human genome by combining a wide range of genomic annotations derived from various sources, including conservation across species, gene and transcript models, protein language models, etc. We categorize CADD-1.7 as an alignment-based model due to its reliance on PhyloP conservation scores.

### Multispecies data

For multispecies training of large language models, we considered the 241 mammalian genomes of the Zoonomia project (Zoonomia Consortium, 2020). The Zoonomia dataset is particularly attractive as it also provides a Phylop-241way model, yielding an evolutionary-based conservation score computed on whole genome alignment derived using one of the most recent aligners named Progressive Cactus (Armstrong *et al*., 2020).

Training data for the 3’UTR-specific models was prepared as follows. 3’UTR coordinates for 18,134 protein-coding genes of the human genome were extracted using BioMart (https://www.ensembl.org/biomart/martview/), considering the Ensembl canonical transcript for each gene. As a rough estimation of 3’UTR coordinates in non-human genomes, we considered sequence segments downstream of the stop codon, with the length corresponding to the 3’UTR length of the respective human transcript (Fig. S1). Sequences mapped to the negative (reverse) strand of the human DNA, including human transcripts on the negative strand, were reverse complemented to match the RNA content. The positions of the stop codons with respect to the human genome and the strand information were derived from the Zoonomia Progressive Cactus alignment.

### Recognition of RBP binding motifs

The positive set of RNA binding protein (RBP) motifs was built based on the consensus of two experiments that utilized distinct methodologies: RNA Bind-n-Seq (RBNS) and enhanced crosslinking and immunoprecipitation (eCLIP). Specifically, we considered RBNS-detected 5-mers (Dominguez *et al*., 2018) overlapping with eCLIP sites (Van Nostrand *et al*., 2020). Similarly to (Griesemer *et al*., 2021), we included in the positive set only 5-mers with strong experimental evidence (Stepwise_R-1>0.1), which resulted in 153 unique motifs. All other 5-mers and 5-mers not overlapping with eCLIP sites were assigned to the negative set. To reduce the computational burden, this analysis was performed on 25% of human 3’UTR sequences. In total, we considered 723,141 and 7,169,129 unique motif hits in the positive and negative sets, correspondingly.

For each motif, we computed a distance-based conservation score, defined as the number of genomes in the Zoonomia whole genome alignment that contain a given motif within a specific window of W basepairs centered at the motif position in the human sequence. Motif hits in non-human genomes were assigned to the nearest positions of the same motif in the human genome. Overlapping windows were not permitted: for each motif hit in the human genome, the conservation window could not extend beyond the next position of the same motif in the human sequence. The mobility of each motif was computed as the difference between conservation scores for W=W*≠0 and W=0.

The enrichment for functional motifs was computed as the odds ratio for putative functional elements contained in the top 10th percentile and in the bottom 10th percentile of the per-base reference allele probability (p_ref_) averaged over the motif positions (n=1..5). This probability was derived from language model predictions, as described in the next section. We consider a higher odds ratio as an indicator of a better model performance. Assuming that a more functional element is more conservative, it can be expected to occur repetitively within a given sequence context in the dataset. Such elements should then be easier to reconstruct for models which effectively leverage the sequence context. Hence, the top 10th percentile of p_ref_ should contain a larger fraction of functional variants than the bottom 10th percentile, leading to a higher odds ratio. For the PhyloP models, the odds ratio was computed based on the conservation score delivered by the model. Odds ratios computed on the bottom 10% and the top 10% of mobility distribution are shown for different W* on Fig. S2.

For the DNABERT-2 models, p_ref_ could not be computed since the variable token length in the BPE encoding scheme impedes separation of contributions from individual positions along the sequence. For CADD-1.7, the enrichment for functional motifs could not be estimated as the model does not provide a way to construct a consistent per-motif score.

### Prioritization of functional variants

We used four different sources to construct four positive sets of functional variants. As putative functional variants, we first considered SNPs with (likely) pathogenic annotations from ClinVar v. 2023.10.07 (Landrum *et al*., 2018), dropping variants with *no_assertion* or *no_interpretation* annotations. However, as a high conservation score from PhyloP often serves as a criterion to include non-coding variants to ClinVar, the estimation of generalization performance of the PhyloP-100way model based on ClinVar variants can be compromised. Hence, we also considered rare SNPs with allelic count (AC) of 1 from gnomAD v. 3.1.2 (Chen *et al*., 2022) as an alternative set of putative functional variants. As the third set, we utilized SNPs associated with gene expression, known as expression quantitative trait loci (eQTLs), extracted from the eQTL-SuSiE fine-mapping credible sets (Kerimov *et al*., 2021) with a stringent P-value threshold of <10^-12^. Finally, we selected 3’UTR-specific ‘proxy-deleterious’ variants used for CADD-1.7 training (Schubach *et al*., 2024). This is a set of simulated variants that do not naturally occur in primates and might thus be enriched for functional mutations. In total, we retained 215 ClinVar, 2,788,551 gnomAD, 21,448 eQTL, and 153,856 CADD SNPs.

For each positive set out of ClinVar, gnomAD, and eQTL data, we built a set of negative variants using 10,000 randomly chosen SNPs with gnomAD population allele frequency (AF) above 5%. These SNPs were selected to ensure no overlap with any positive variant. As a matched set for the ‘proxy-deleterious’ CADD variants, we selected the corresponding number of 3’UTR-specific ‘proxy-neutral’ variants, also used in CADD training. These are variants that naturally occur in primates and are thus more likely to be benign. To reduce the computational burden, we used at most 10,000 positive or negative variants in each dataset.

We then calculated a set of scores that could be used as an indicator of a variant’s functional significance. We call them *zero-shot* scores, aligning with the machine learning paradigm where a pre-trained model is evaluated on classes that were not used at training. Three of these scores were computed based on model predictions at the variant position: the reference allele probability (p_ref_), the logarithm of inverse alternative allele probability (log(p^-1^_alt_)), and the logarithmic ratio of these two probabilities (log(p_ref_/p_alt_)). Another six scores were computed based on various distance metrics between model embeddings for reference and alternative sequences (l1, l2, dot product, and cosine similarity) and based on model losses (loss on the alternative sequence and the difference between the alternative and reference losses). For each functionality score and each SNP dataset, we also computed the ROC AUC measure. These ROC AUC scores are reported alongside with their bootstrap-estimated 95% confidence intervals in Table S2. We considered two models equivalent when their 95% confidence intervals for ROC AUC overlap.

To derive per-nucleotide probabilities for DNABERT, NT-MS-v2-250M, and the state space architectures, we consecutively masked each 6-mer token in the sequence. The sum over final layer softmax probabilities for all tokens carrying a given base in a given position within the masked 6-mer was then used as a per-base prediction. Since DNABERT is based on overlapping tokenization, we masked a contiguous span of 6 tokens instead of a single k-mer. This technique was previously used in (Gankin *et al*., 2023; Karollus *et al*., 2023). For the 3’UTR-specific models, we used RNA instead of DNA sequences when computing embeddings, in agreement with the Multispecies dataset preparation and model training procedure.

In addition to zero-shot scores, we trained task-specific supervised models for variant effect prediction based on language model embeddings as input. For each variant, the embeddings were generated on 1024-bp long sequences centered around the variant position. The embeddings for the reference and alternative alleles were concatenated, resulting in a 2x longer vector. We used a fully connected multilayer perceptron (MLP) evaluated in a nested Cross-Validation (CV) fashion, using the inner 5-fold CV for the hyperparameter search and the outer 10-fold CV for the estimation of model generalization performance. The MLP was trained for 300 epochs with learning rate of 1e-4 and batch size of 1024 using the AdamW optimizer (Loshchilov and Hutter, 2019). The hyperparameter search for the number of layers, dropout probability, and weight decay was performed by running a Tree-structured Parzen estimator solver (Bergstra *et al*., 2011) 150 times. To reduce the computational burden, we retained the hyperparameters determined for the initial (first) outer CV split. The MLP was trained separately for each language model and each SNP dataset, resulting in a total of 24 models. The resulting ROC AUC scores on the hold out data and their bootstrap-estimated 95% confidence intervals are reported in Table S3. We considered two models equivalent when their 95% confidence intervals for ROC AUC overlap.

### Prediction of MPRA expression levels and mRNA half-life

To predict expression levels in massively parallel reporter assays (MPRA) and to estimate mRNA half-life, we used Ridge Regression and Support Vector Regression (SVR) models trained on language model embeddings.

We also trained baseline Ridge and SVR models using the features suggested in the original study. Specifically, for the MPRA data from (Griesemer *et al*., 2021) we utilized the following attributes of the *minimal model* which was proposed to predict a binary label indicating attenuation or increase in expression due to a specific variant: nucleotide percentage, dinucleotide percentages, exact dinucleotide counts, maximum homopolymer length, maximum dinucleotide length across all bases, and a measure of sequence uniformity. For the MPRA data from (Siegel *et al*., 2022), the baseline model was trained using 5-mer sequence embeddings since these features led to the best predictions in the original study.

We considered the mRNA steady state measured in six cell lines as prediction targets for the (Griesemer *et al*., 2021) data. For the (Siegel *et al*., 2022) data, we predicted the mRNA steady state and mRNA stability assessed in the initial study for two distinct cell lines. Similarly to the original studies, we predicted the expression levels independently for oligo sequences carrying the reference and alternative alleles. The length of oligo sequences used to generate embeddings was at most 101bp for the (Griesemer *et al*., 2021) data and 160bp for the (Siegel *et al*., 2022) data, in agreement with the original studies.

For convenience, we excluded oligos with alternative allelic background from the (Griesemer *et al*., 2021) data. In total, we generated embeddings for 15,970 oligo sequences from the (Griesemer *et al*., 2021) experiment and 10,609 oligo sequences from the (Siegel *et al*., 2022) experiment, on average per cell type and per prediction target.

For the 3’UTR-specific models, we used RNA instead of DNA sequences when computing embeddings, in agreement with the Multispecies dataset preparation and model training procedure.

The generalization performance was estimated using LeaveOneGroupOut CV. The grouping criteria followed the conventions of the original study: groups for the (Griesemer *et al*., 2021) data were composed of oligos derived from the same gene, groups for the (Siegel *et al*., 2022) data matched chromosomes, and groups for the (Agarwal and Kelley, 2022) data aligned with the original folds. The prediction quality was measured with the Pearson correlation coefficient r, computed between the predictions gathered from all test folds and the values measured in the experiment. We considered two models equivalent when their 95% confidence intervals for Pearson r overlap. The 95% confidence interval was computed using Fisher transformation.

The Ridge regression and SVR models were evaluated in a nested CV fashion, using the inner 5-fold group-based CV for the hyperparameter search and the outer LeaveOneGroupOut CV for the estimation of model generalization performance. For SVR, we retained the hyperparameters determined for the initial (first) outer CV split to reduce the computational burden. The SVR hyperparameters (regularization strength *C*, RBF kernel coefficient *γ*, and epsilon-tube width *ε*) were searched on a logarithmic scale by running a Tree-structured Parzen estimator solver (Bergstra *et al*., 2011) 300 times.

The CADD-1.7 model was excluded from evaluation in embeddings-based tasks since it does not provide a way to generate sequence embeddings.

## Results

We first assessed the ability of each model to detect RBP binding sites represented by 153 unique 5-mers overlapping with eCLIP positions. This task can be particularly challenging for alignment-based models since regulatory elements might exhibit certain mobility (Gankin *et al*., 2023; Karollus *et al*., 2023): in the context of whole genome alignment, the exact position of a given regulatory element may vary across different genomes and not necessarily match its position in the human sequence.

For each model, we estimated the enrichment of potential functional motifs using odds ratios computed based on the reference allele probability p_ref_, as defined in the **Methods** section. Fig. 1 illustrates the resulting performance for the conservation window of W*=5000bp, for which the odds ratios stabilize (Fig. S2.). The best performance is delivered by the StateSpace-SA model (Fig. 1a), followed by StateSpace and DNABERT-3UTR. It is important to note that the 3’UTR-specific models DNABERT-3UTR and NTv2-250M-3UTR largely outperform their whole genome versions. The highly skewed mobility distribution (Fig. 1b) indicates a large fraction of stationary and weakly mobile motifs in the human genome. When splitting motifs according to their mobility, all models recognize weakly mobile motifs better compared to the high-mobility group (Fig. 1c,d). The PhyloP models perform the best in the low-mobility category (bottom 10% mobility, Fig. 1c) while StateSpace-SA ranks first in the high-mobility category (top 10% mobility, Fig. 1d).

**Fig. 1.**
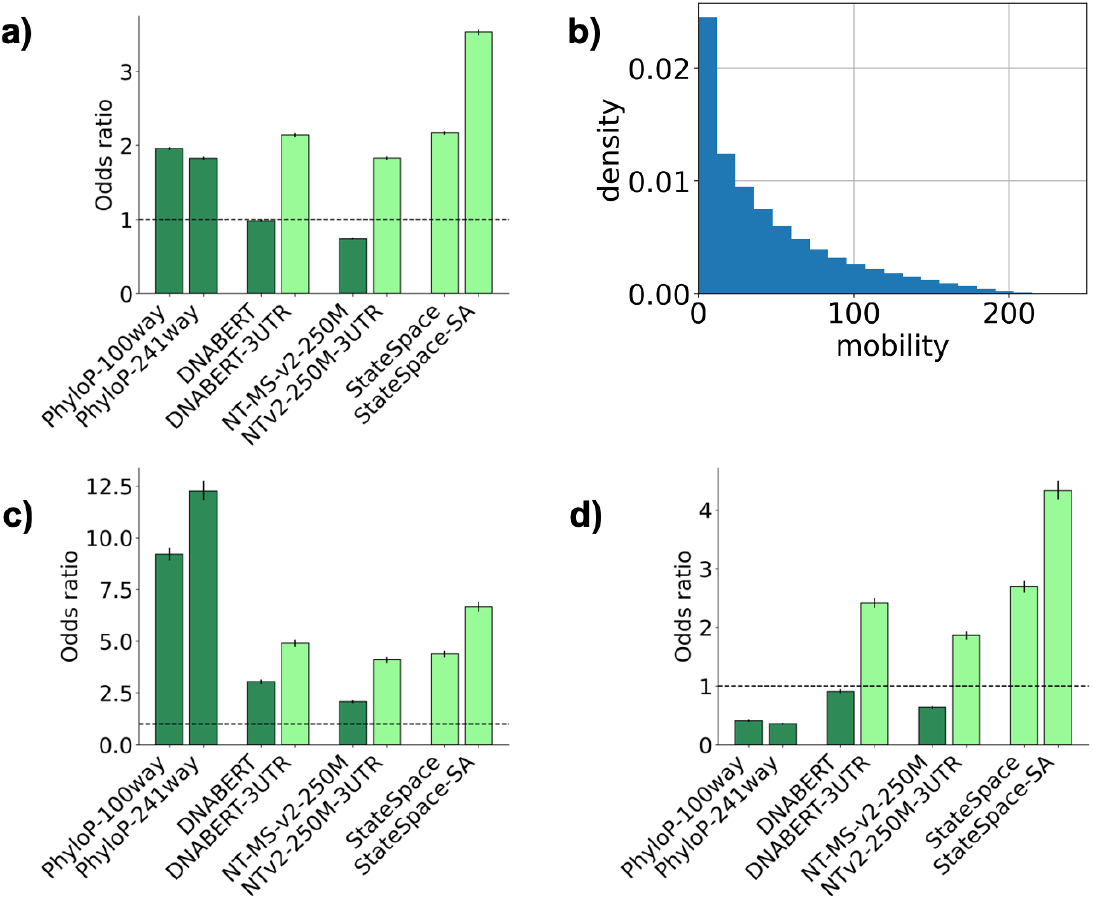
(a) Enrichment for all putative functional motifs, (b) mobility distribution for putative functional motifs, (c) enrichment for putative functional motifs within the bottom 10% mobility (W*=5000bp), (d) enrichment for putative functional motifs within the top 10% mobility (W*=5000bp). The error bars show the 95% confidence intervals.

We also highlight that the model performance reaches stability at higher values of the conservation window width W* (Fig. S2.). A greater W* likely results in a more accurate estimation of mobility by considering more distant hits.

We next estimated the ability of the analyzed models to prioritize functional variants in 3’UTR sequences. To this end, we first computed Receiver Operating Characteristic Area Under the Curve (ROC AUC) on 9 zero-shot scores for the ClinVar, gnomAD, eQTL, and CADD datasets (Table S2). In this setting, the language models specifically trained on 3’UTR sequences either outperformed the genome-wide models (ClinVar, gnomAD, eQTL) or demonstrated equivalent results (CADD), with the highest score achieved on ClinVar (AUC=0.765, NTv2-250M-3UTR), followed by CADD (AUC=0.666, NT-MS-v2-250M), gnomAD (AUC=0.593, StateSpace), and eQTL (AUC=0.555, StateSpace). Even in this simple zero-shot scenario the 3’UTR-specific language models performed better or equivalently to PhyloP-241way, also trained on the Zoonomia dataset (Table S3).

We then trained a MLP classifier to predict functional variants based on language model embeddings. The resulting ROC curves are shown in Fig. 2, with the AUC scores summarized in Table S3. Compared to zero-shot prediction, most of the language models improved in performance for all the four datasets. It is also of note that the DNABERT-2 and the DNABERT2-ZOO models demonstrated equivalent results.

**Fig. 2.**
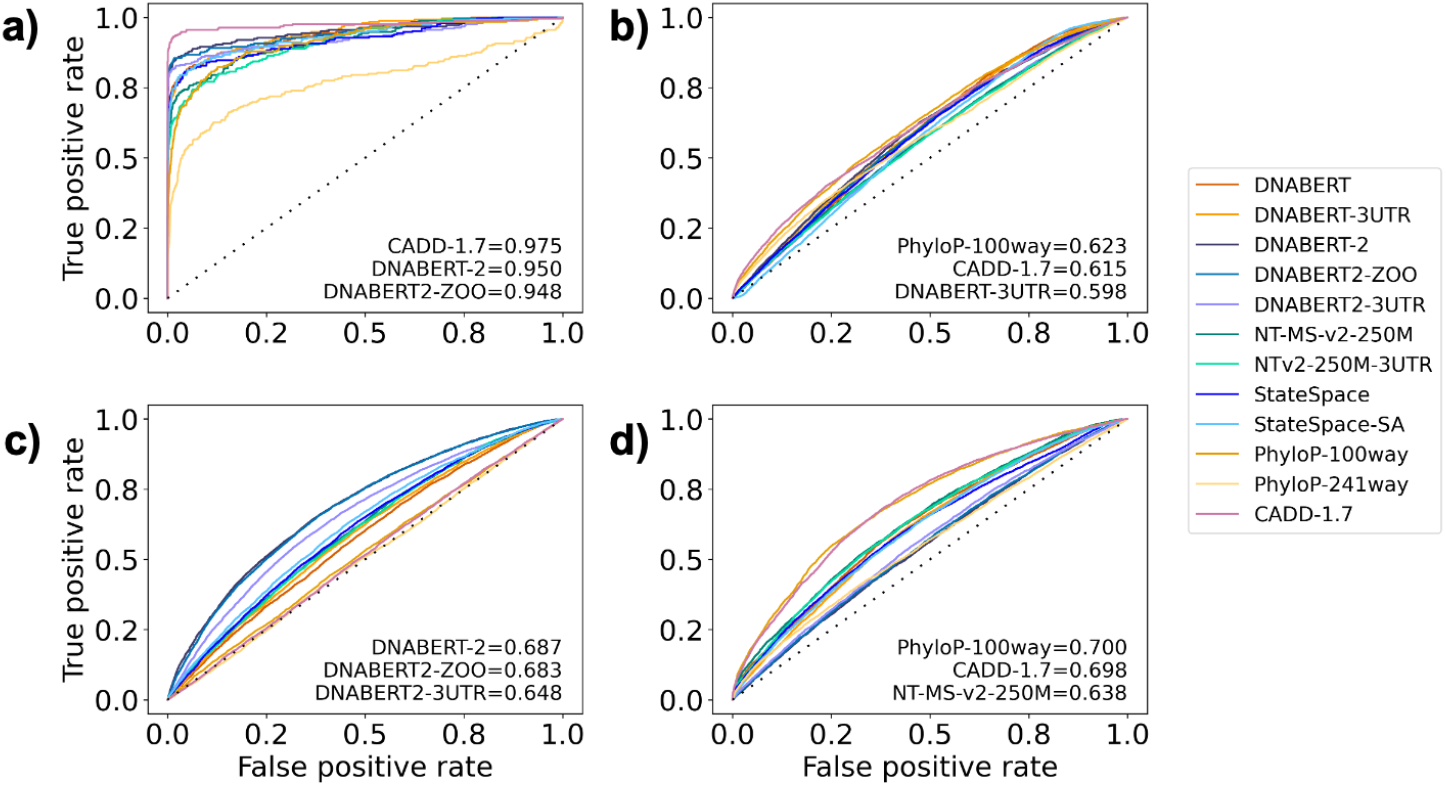
ROC curves resulting from prediction of functional variants on (a) ClinVar, (b) gnomAD, (c) eQTL, and (d) CADD data using language model embeddings and PhyloP scores.The ROC AUC scores for the three best models are shown.

We further explored the potential of language model embeddings by employing them for two downstream tasks, namely for predicting reporter expression in two MPRA studies on 3’UTR variants and for predicting mRNA half-life. MPRA is a powerful experimental technique to explore the regulatory impact of variants. The alleles associated with the analyzed variants are first seeded into short oligo-nucleotide sequences which are placed next to the reporter gene on a plasmid. By assessing reporter expression across different oligo sequences, the impact of the analyzed variants (alleles) on gene expression can be estimated.

We first applied Ridge regression to predict MPRA reporter expression from the embeddings. MPRA data was measured in six human cell lines for several thousand variants associated with human disease and evolutionary selection (Griesemer *et al*., 2021). The Pearson correlation coefficient r between expression predictions and the hold out data is shown in Table 1.

**Table 1.** Pearson r correlation coefficient between Ridge-based predictions from sequence embeddings and ground truth MPRA expression from (Griesemer *et al*., 2021).

|  | HEK293FT | HMEC | HEPG2 | GM12878 | K562 | SKNSH |
| --- | --- | --- | --- | --- | --- | --- |
| **DNABERT** | 0.17±0.02 | 0.38±0.01 | 0.27±0.01 | 0.28±0.01 | 0.23±0.01 | 0.22±0.01 |
| **DNABERT-3UTR** | **0.33±0.01** | **0.51±0.01** | 0.36±0.01 | 0.37±0.01 | **0.37±0.01** | **0.35±0.01** |
| **DNABERT-2** | 0.10±0.02 | 0.31±0.01 | 0.13±0.02 | 0.11±0.02 | 0.18±0.01 | 0.15±0.02 |
| **DNABERT2-ZOO** | 0.08±0.02 | 0.31±0.01 | 0.14±0.02 | 0.12±0.02 | 0.15±0.02 | 0.11±0.02 |
| **DNABERT2-3UTR** | 0.16±0.02 | 0.42±0.01 | 0.30±0.01 | 0.30±0.01 | 0.25±0.01 | 0.20±0.01 |
| **NT-MS-v2-250M** | 0.14±0.02 | 0.31±0.01 | 0.18±0.02 | 0.15±0.02 | 0.20±0.01 | 0.18±0.02 |
| **NTv2-250M-3UTR** | 0.25±0.01 | 0.45±0.01 | 0.28±0.01 | 0.28±0.01 | 0.29±0.01 | 0.27±0.01 |
| **StateSpace** | 0.30±0.01 | 0.49±0.01 | 0.38±0.01 | 0.39±0.01 | 0.33±0.01 | 0.32±0.01 |
| **StateSpace-SA** | **0.32±0.01** | **0.51±0.01** | **0.41±0.01** | **0.42±0.01** | 0.35±0.01 | **0.33±0.01** |
| **Griesemer et al., 2021** | 0.22±0.01 | 0.42±0.01 | 0.25±0.01 | 0.24±0.01 | 0.28±0.01 | 0.25±0.01 |

The StateSpace-SA and DNABERT-3UTR models yield the highest correlation. For the DNABERT, DNABERT-2, and NT-MS-v2-250M architectures, region-specific training leads to a better performance compared to using whole genome data. As for the previous task, the whole genome Zoonomia-based DNABERT2-ZOO model performs on par with the original DNABERT-2 model.

Embeddings generated by distinct model architectures encode data differently. For some of them, Ridge regression may lack the complexity required to reveal intricate patterns relevant to gene expression, which might hinder an objective comparison. We therefore trained a more sophisticated SVR regressor with a RBF kernel. SVR greatly improves the prediction score for all the models (Table S4), with embeddings from the StateSpace-SA model leading to the best or equivalent performance across all cell lines.

We then evaluated the potential utility of language model embeddings in predicting mRNA steady-state levels and stability in Jurkat and BeasB2 cells measured in the second MPRA experiment (Siegel *et al*., 2022). The prediction results are shown in Table S5 and Table S6 for Ridge and SVR regressors correspondingly. In this task, the DNABERT-3UTR model achieves the best or equivalent performance. However, the difference between Ridge and SVR results is less pronounced. We observed that training on this dataset required stronger regularization (smaller SVR parameter *C*), which may level out the performance difference between simple and more complex machine learning techniques.

Finally, we used embeddings generated by language models to predict mRNA half-life derived by (Agarwal and Kelley, 2022). In that study, the consensus mRNA half-life was obtained as the 1st principle component of a *sample x gene* matrix composed of half-life measurements from 39 human samples, spanning different cell types and measurement techniques. We first predicted the half-life from 3’UTR sequences only, using various 3’UTR embeddings (Fig. 3a). In this task, the NTv2-250M-3UTR, StateSpace, and StateSpace-SA models demonstrated the best performance. As for the previous task, the results improved under SVR (Table S7). Among the language models, DNABERT and DNABERT-3UTR led to the least accurate predictions, likely due to their limited field of view (512bp). Again, the DNABERT-2 and DNABERT2-ZOO models showed equivalent results. Although the k-mer embeddings performed the worst when using ridge regression, under SVR they performed equivalently to the language models.

**Fig. 3.**
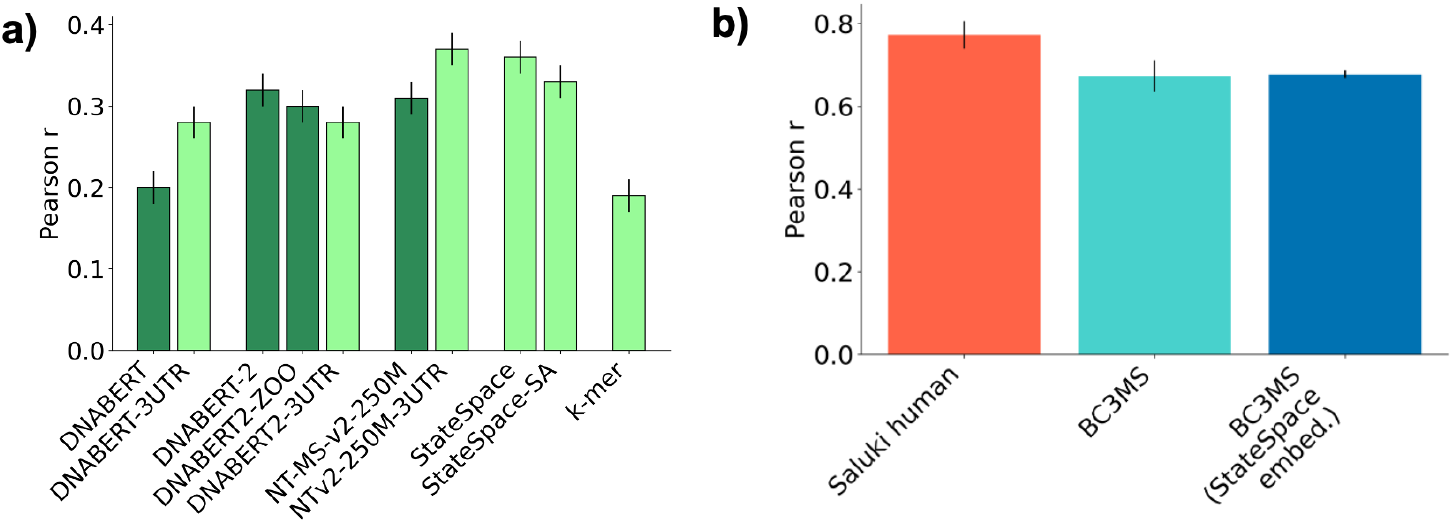
(a) Pearson r correlation coefficient between mRNA half-life prediction and ground truth data from (Agarwal and Kelley, 2022), using Ridge regression applied to different 3'UTR embeddings, (b) Pearson r correlation coefficient for mRNA half-life prediction with the BC3MS model based on different 3'UTR embeddings and the Saluki model. The performance of the BC3MS model with StateSpace embeddings is reported based on the SVR results. The performance of the original BC3MS (k-mer) and Saluki models is reported based on the predictions from the original study (Agarwal and Kelley, 2022). The error bars show the 95% confidence intervals.

We then predicted mRNA half-life using the BC3MS model (Agarwal and Kelley, 2022). In the original study, this model showed the best performance among all models utilizing manually crafted sequence-based features. The original BC3MS architecture relies on basic mRNA features (length and G/C content of 5’UTR, ORF, and 3’UTR; intron length; ORF exon junction density), mRNA codon counts, 3’UTR *k*-mers (*k*=1..7), miRNA target repression and SeqWeaver RBP binding prediction. To generate mRNA half-life predictions, we replaced *k*-mer encodings for 3’UTR sequences with embeddings from the StateSpace model. As follows from Fig. 3b, both encodings demonstrate equivalent performance. However, the resulting Pearson r coefficient is notably lower than that achieved by the deep Saluki model (Agarwal and Kelley, 2022) trained based on the entire mRNA sequence.

## Discussion

In this work, we considered a range of 3’UTR-specific tasks to compare general-purpose genomic foundation models with multispecies language models specifically trained on 3’UTR sequences and conservation-based PhyloP models. In most of the tasks, language models specifically trained on 3’UTR sequences outperformed their whole genome counterparts.

We first proceeded with identification of RBP binding motifs. In this task, the StateSpace-SA model demonstrated superior performance. It is also of note that the language models were largely outperformed by the PhyloP models when identifying stationary or weakly mobile RBP motifs. In contrast, the language models demonstrated their ability to leverage the specific sequencing context in an alignment-free manner by outperforming the alignment-based PhyloP models on highly mobile motifs. The PhyloP models can be expected to fail on this category since they treat the alignment column-wise and a minor motif displacement will lead to a decreased conservation score.

Our mobility analysis revealed a large fraction of stationary and weakly mobile functional motifs. A possible reason for this is the relatively close evolutionary distances between the Zoonomia species: all of these species are mammals spanning around 110 million years of evolution. Shorter evolutionary distances may explain little variations in the motif position across different species in the alignment.

We then proceeded with variant effect prediction. We first observed that the language models exhibit certain predictive capacity by delivering meaningful zero-shot scores, with the 3’UTR state space models outperforming their genome-wide counterparts. Notably, on the ClinVar and eQTL datasets the single-position scores were outperformed by the scores based on embeddings and losses. Perturbing a single (variant) position in a sequence results in an altered predicted probability for all associated loci. These alterations are then accumulated in losses and embeddings which should therefore provide a stronger signal than a single score at the variant position. Compared to p_ref_, the scores derived from embeddings or loss differences also account for the alternative sequence, which should further improve the signal-to-noise ratio. The weak signal from p_ref_ might actually be the reason why the PhyloP models outperform the language-based models on stationary or weakly mobile RBP motifs.

Task-specific supervised learning based on sequence embeddings significantly improved variant classification for 3 out of 4 datasets, suggesting that the operations applied on embeddings to compute zero-shot scores might not be complex enough to capture a biologically meaningful distance between the reference and alternative sequences. This improvement stands in agreement with previous observations (Dalla-Torre *et al*., 2023). Note that the language models trained on the Zoonomia species showed better results than the PhyloP-241way model built on the same dataset.

The superiority of region-specific language models was further corroborated when predicting MPRA expression measured in the experiments of (Griesemer *et al*., 2021) and (Siegel *et al*., 2022). In this task, all region-specific models outperformed their genome-wide counterparts, with the StateSpace-SA and DNABERT-3UTR models scoring the best.

The region-specific NTv2-250M-3UTR and state space models also performed the best when predicting mRNA half-life based on 3’UTR embeddings only. In this task they outperformed *k*-mer (*k*=1..7) embeddings used to encode 3’UTR sequences in the original study (Agarwal and Kelley, 2022). However, addition of other features (BC3MS model) completely annihilated the performance difference, probably due to these features also including 3’UTR-specific information, such as G/C content and predicted average binding score of human RBPs, alongside with the information from the remaining mRNA. Notably, using language model embeddings as an input to the BC3MS model did not reach the performance of the deep Saluki model (Agarwal and Kelley, 2022). This suggests that the superior performance of the full-sequence deep Saluki model can not be attributed to enhanced encoding of the 3’UTR sequence alone, but rather to inferring additional information from other segments of the mRNA sequence.

In all tasks except variant effect prediction, the models trained specifically on 3’UTR regions outperformed their genome-wide counterparts. This notable performance difference can not be solely explained by different sets of species used in training, as evidenced by the DNABERT-2 and DNABERT2-ZOO models showing equivalent performance across all the tasks. We hypothesize that the established foundation models trained on whole genome data might confuse contextual information and dependencies arising from different genomic regions. For example, it was pointed out (Dalla-Torre *et al*., 2023) that the NT models might encounter challenges in recognizing 3’UTR sequences. We also note that our multispecies dataset was constructed in such a way that 3’UTR sequences of human genes located on the negative strand, as well as sequences from other species aligned to the negative strand of the human genome, were reverse complemented to align with mRNA content. This transforms the multispecies dataset from completely species-agnostic DNA-based into human RNA-centric and should help the models to discover common patterns in 3’UTR regions across the species in the dataset. To our knowledge, we are the first to incorporate the whole genome alignment strand information into a language model.

We also note that across all the tasks, the StateSpace-SA model outperformed or performed on par with its species-agnostic version. This result is consistent with the work of (Gankin *et al*., 2023; Karollus *et al*., 2023) who showed that species-awareness leads to an improved performance.

The observed performance difference between 3’UTR-specific models is apparently related to their architectures and is a subject of a future study.

Finally, we would like to highlight that the proposed set of tasks focusing on 3’UTRs expands the range of applications of language models in the field of human RNA research. This set of tasks can also serve as a benchmark for future model development.

## Conclusion

We demonstrated that language models trained on 3’UTR sequences from a large set of mammalian genomes outperform existing human-based and multispecies foundation models on three out of four 3’UTR-specific tasks. Our results highlight the importance of evaluating language models in a region-specific manner. Additionally, the proposed set of 3’UTR-specific tasks can be used as a benchmark for future model development.

## Acknowledgement

We would like to thank Julien Gagneur for an insightful discussion and the provided computational resources of the Computational Molecular Medicine lab that supported the start of this work.

## Data and code availability

The pre-trained weights and the model implementation of the 6-mer DNABERT model were downloaded from the official GitHub repository: https://github.com/jerryji1993/DNABERT. The pre-trained weights and the model implementation of DNABERT-2 were downloaded from the official HuggingFace page: https://huggingface.co/zhihan1996/DNABERT-2-117M. The pre-trained weights and the model implementation of the NT-MS-v2-250M multispecies model were downloaded from the official HuggingFace page: https://huggingface.co/InstaDeepAI/nucleotide-transformer-v2-250m-multi-species. PhyloP-100way conservation scores were downloaded from the UCSC ftp server: http://hgdownload.soe.ucsc.edu/goldenPath/hg38/phyloP100way/. PhyloP241-way conservation scores and Zoonomia Progressive Cactus alignment were downloaded from the UC Santa Cruz Computational Genomics Lab web page: https://cglgenomics.ucsc.edu/data/cactus/. ClinVar data was downloaded from the NCBI ftp server: https://ftp.ncbi.nlm.nih.gov/pub/clinvar/. GnomAD data was derived from the official web page: https://gnomad.broadinstitute.org/. eQTL-SuSiE data was downloaded from the EBI ftp server: http://ftp.ebi.ac.uk/pub/databases/spot/eQTL/susie/. CADD training data was derived from the official CADD website: https://cadd.bihealth.org/. The implementation of the state space architecture was adopted from https://github.com/DennisGankin/species-aware-DNA-LM.

The scripts relevant to this study can be found at: https://github.com/sergeyvilov/investigating-foundation-models-3utr.

