## Supplementary Data for "Investigating the performance of foundation models on human 3’UTR sequences"

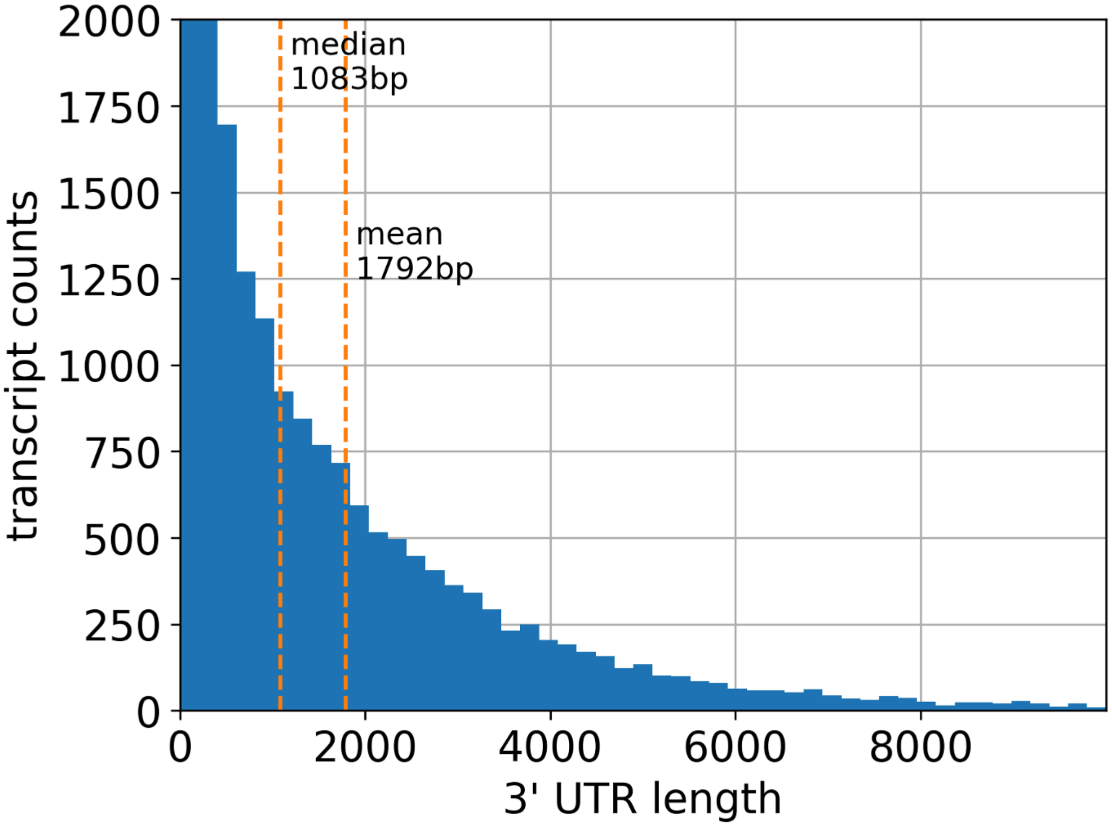


Fig. S1. Distribution of 3’UTR length for 18,134 transcripts of the human genome.

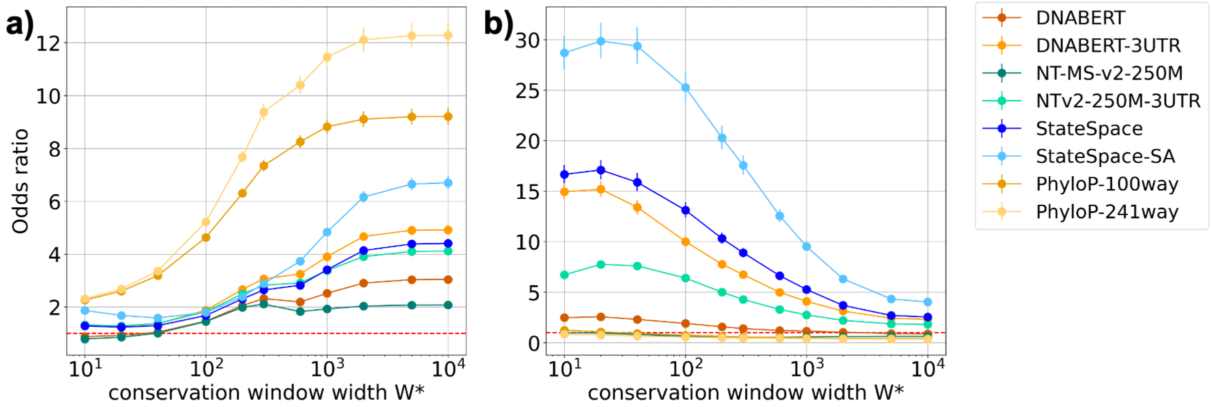


Fig. S2. Enrichment for putative functional motifs as a function of the conservation window width W*: (a) within the bottom 10% mobility, (b) within the top 10% mobility.

| Model | Effective batch size | Max. learning rate | Weight decay | Total training time |
|---|---|---|---|---|
| **DNABERT2-ZOO** | 4480 | 5e-4 | 1e-5 | 168.7 h |
| **DNABERT-3UTR** | 1920 | 4e-4 | 1e-2 | 5.3 h |
| **DNABERT2-3UTR** | 4480 | 5e-4 | 1e-5 | 1.3 h |
| **NTv2-250M-3UTR** | 480 | 1e-4 | 0 | 13.7 h |
| **StateSpace** | 2560 | 1e-4 | 0 | 1 h |
| **StateSpace-SA** | 2560 | 1e-4 | 0 | 1 h |

Table S1. Training parameters for all the models when trained on 10 NVIDIA A100 80G GPUs in parallel for 2 epochs. Learning rate schedules for DNABERT-3UTR, DNABERT2-3UTR, and NTv2-250M-3UTR are applied in agreement with the original model publication.

| dataset | score | DNABERT | DNABERT -3UTR | DNABERT-2 | DNABERT2 -ZOO | DNABERT2 -3UTR | NT-MS-v2 -250M | NTv2-250M -3UTR | StateSpace | StateSpace -SA |
|---|---|---|---|---|---|---|---|---|---|---|
| clin var | $p_{ref}$ | 0.518±0.032 | 0.526±0.035 | - | - | - | 0.514±0.033 | 0.513±0.030 | 0.532±0.032 | 0.520±0.037 |
| | $\log(p^{-1}_{alt})$ | 0.536±0.032 | 0.543±0.032 | - | - | - | 0.589±0.033 | 0.527±0.031 | 0.529±0.031 | 0.526±0.034 |
| | $\log(p_{ref}/p_{alt})$ | 0.506±0.033 | 0.505±0.034 | - | - | - | 0.551±0.033 | 0.521±0.029 | 0.510±0.033 | 0.503±0.032 |
| | l1 | 0.609±0.030 | 0.555±0.031 | 0.628±0.038 | 0.521±0.037 | 0.588±0.034 | **0.698±0.033** | 0.561±0.028 | 0.541±0.039 | 0.520±0.034 |
| | l2 | 0.606±0.030 | 0.555±0.030 | 0.640±0.035 | 0.526±0.038 | 0.588±0.035 | **0.693±0.033** | 0.559±0.029 | 0.541±0.035 | 0.524±0.034 |
| | dot | **0.692±0.031** | 0.605±0.043 | 0.536±0.023 | 0.641±0.029 | 0.598±0.030 | 0.583±0.032 | **0.741±0.032** | 0.605±0.029 | 0.615±0.027 |
| | cosine | 0.556±0.032 | 0.522±0.030 | 0.636±0.034 | 0.515±0.036 | 0.591±0.037 | **0.684±0.037** | 0.583±0.035 | 0.508±0.038 | 0.519±0.038 |
| | $loss_{alt}$ | 0.523±0.033 | 0.533±0.035 | 0.641±0.031 | 0.670±0.036 | 0.541±0.037 | 0.638±0.032 | 0.511±0.036 | 0.635±0.031 | 0.622±0.035 |
| | $loss_{alt}$-$loss_{ref}$ | 0.524±0.035 | 0.505±0.033 | 0.536±0.033 | 0.506±0.033 | 0.510±0.036 | 0.512±0.038 | 0.523±0.036 | 0.502±0.035 | 0.501±0.035 |
| gnom AD | $p_{ref}$ | 0.571±0.008 | 0.566±0.007 | - | - | - | 0.536±0.008 | 0.538±0.008 | 0.567±0.008 | 0.572±0.008 |
| | $\log(p^{-1}_{alt})$ | **0.581±0.008** | 0.571±0.008 | - | - | - | 0.568±0.008 | 0.563±0.008 | 0.572±0.008 | 0.557±0.008 |
| | $\log(p_{ref}/p_{alt})$ | **0.591±0.007** | **0.584±0.008** | - | - | - | 0.563±0.008 | 0.559±0.008 | **0.584±0.008** | **0.578±0.008** |
| | l1 | 0.518±0.008 | 0.508±0.008 | 0.515±0.008 | 0.518±0.008 | 0.542±0.008 | 0.512±0.008 | 0.511±0.008 | 0.560±0.008 | 0.541±0.008 |
| | l2 | 0.519±0.008 | 0.509±0.008 | 0.514±0.008 | 0.518±0.008 | 0.542±0.008 | 0.511±0.008 | 0.511±0.008 | 0.561±0.008 | 0.542±0.008 |
| | dot | 0.513±0.008 | 0.513±0.008 | 0.536±0.008 | 0.532±0.008 | 0.509±0.008 | 0.520±0.008 | 0.520±0.008 | 0.521±0.007 | 0.522±0.008 |
| | cosine | 0.523±0.008 | 0.504±0.008 | 0.526±0.008 | 0.524±0.008 | 0.543±0.008 | 0.509±0.008 | 0.518±0.008 | 0.565±0.008 | 0.541±0.008 |
| | $loss_{alt}$ | 0.516±0.008 | 0.514±0.008 | 0.513±0.008 | 0.503±0.008 | 0.503±0.008 | 0.510±0.007 | 0.522±0.008 | 0.516±0.008 | 0.515±0.008 |
| | $loss_{alt}$-$loss_{ref}$ | 0.501±0.008 | 0.528±0.008 | 0.513±0.008 | 0.506±0.008 | 0.505±0.008 | 0.507±0.008 | 0.512±0.008 | **0.586±0.008** | **0.581±0.008** |
| eQTL | $p_{ref}$ | 0.512±0.008 | 0.509±0.008 | - | - | - | 0.512±0.008 | 0.504±0.008 | 0.509±0.008 | 0.513±0.008 |
| | $\log(p^{-1}_{alt})$ | 0.501±0.008 | 0.502±0.008 | - | - | - | 0.502±0.008 | 0.502±0.008 | 0.501±0.008 | 0.501±0.008 |
| | $\log(p_{ref}/p_{alt})$ | 0.507±0.008 | 0.506±0.008 | - | - | - | 0.505±0.008 | 0.501±0.008 | 0.505±0.008 | 0.508±0.008 |
| | l1 | 0.511±0.008 | 0.509±0.008 | 0.507±0.008 | 0.508±0.008 | 0.506±0.008 | 0.501±0.008 | 0.514±0.008 | 0.508±0.008 | 0.518±0.008 |
| | l2 | 0.510±0.008 | 0.508±0.008 | 0.508±0.008 | 0.508±0.008 | 0.506±0.008 | 0.501±0.008 | 0.514±0.008 | 0.507±0.008 | 0.518±0.008 |
| | dot | 0.524±0.008 | 0.507±0.008 | **0.552±0.008** | 0.535±0.008 | 0.510±0.008 | 0.523±0.008 | 0.522±0.008 | **0.544±0.008** | 0.527±0.008 |
| | cosine | 0.504±0.008 | 0.508±0.008 | 0.511±0.008 | 0.502±0.008 | 0.507±0.008 | 0.504±0.008 | 0.501±0.008 | 0.522±0.008 | 0.521±0.008 |
| | $loss_{alt}$ | 0.509±0.008 | 0.511±0.008 | 0.528±0.008 | 0.525±0.008 | 0.521±0.008 | 0.515±0.008 | 0.536±0.008 | **0.547±0.008** | 0.533±0.008 |
| | $loss_{alt}$-$loss_{ref}$ | 0.502±0.007 | 0.501±0.008 | 0.505±0.008 | 0.501±0.008 | 0.500±0.008 | 0.503±0.007 | 0.507±0.008 | 0.506±0.008 | 0.508±0.008 |
| CADD | $p_{ref}$ | 0.623±0.008 | 0.605±0.008 | - | - | - | 0.638±0.008 | 0.632±0.008 | 0.610±0.008 | 0.583±0.008 |
| | $\log(p^{-1}_{alt})$ | 0.626±0.008 | 0.598±0.007 | - | - | - | 0.648±0.008 | 0.639±0.007 | 0.602±0.008 | 0.566±0.008 |
| | $\log(p_{ref}/p_{alt})$ | 0.633±0.008 | 0.607±0.007 | - | - | - | **0.666±0.007** | **0.654±0.008** | 0.612±0.008 | 0.576±0.008 |
| | l1 | 0.505±0.008 | 0.510±0.008 | 0.508±0.008 | 0.504±0.008 | 0.503±0.008 | 0.504±0.008 | 0.506±0.008 | 0.527±0.008 | 0.522±0.008 |
| | l2 | 0.506±0.008 | 0.511±0.008 | 0.509±0.008 | 0.504±0.008 | 0.503±0.008 | 0.503±0.008 | 0.505±0.008 | 0.527±0.008 | 0.521±0.008 |
| | dot | 0.516±0.008 | 0.509±0.008 | 0.533±0.008 | 0.526±0.007 | 0.501±0.008 | 0.516±0.008 | 0.514±0.008 | 0.524±0.008 | 0.521±0.008 |
| | cosine | 0.501±0.008 | 0.508±0.008 | 0.504±0.008 | 0.503±0.008 | 0.503±0.008 | 0.501±0.008 | 0.509±0.008 | 0.532±0.008 | 0.521±0.008 |
| | $loss_{alt}$ | 0.513±0.008 | 0.511±0.008 | 0.509±0.008 | 0.517±0.008 | 0.510±0.008 | 0.521±0.008 | 0.521±0.008 | 0.519±0.008 | 0.518±0.008 |
| | $loss_{alt}$-$loss_{ref}$ | 0.504±0.008 | 0.524±0.008 | 0.522±0.008 | 0.506±0.008 | 0.505±0.008 | 0.504±0.008 | 0.522±0.008 | 0.599±0.008 | 0.570±0.008 |

Table S2. ROC AUC scores for ClinVar, gnomAD, eQTL, and CADD data computed based on zero-shot functionality scores for all models.

| Model | ClinVar | gnomAD | eQTL | CADD |
|---|---|---|---|---|
| **DNABERT** | **0.940±0.018** | 0.594±0.008 | 0.573±0.008 | 0.623±0.008 |
| **DNABERT-3UTR** | **0.945±0.017** | 0.598±0.008 | 0.586±0.008 | 0.612±0.007 |
| **DNABERT-2** | **0.950±0.022** | 0.597±0.007 | **0.687±0.007** | 0.549±0.008 |
| **DNABERT2-ZOO** | **0.948±0.018** | 0.588±0.008 | **0.683±0.007** | 0.553±0.008 |
| **DNABERT2-3UTR** | **0.928±0.023** | 0.562±0.008 | 0.648±0.008 | 0.563±0.008 |
| **NT-MS-v2-250M** | **0.918±0.021** | 0.563±0.008 | 0.599±0.008 | 0.638±0.008 |
| **NTv2-250M-3UTR** | 0.912±0.023 | 0.560±0.008 | 0.597±0.008 | 0.632±0.008 |
| **StateSpace** | **0.922±0.024** | 0.586±0.008 | 0.603±0.008 | 0.610±0.008 |
| **StateSpace-SA** | **0.933±0.020** | 0.572±0.008 | 0.616±0.008 | 0.613±0.007 |
| **PhyloP-100way** | 0.915±0.022 | **0.623±0.008** | 0.519±0.008 | **0.700±0.007** |
| **PhyloP-241way** | 0.774±0.038 | 0.574±0.008 | 0.501±0.008 | 0.557±0.008 |
| **CADD-1.7** | **0.975±0.018** | **0.615±0.008** | 0.510±0.008 | **0.698±0.007** |

Table S3. ROC AUC scores from prediction of functional variants on ClinVar, gnomAD, eQTL, and CADD data using language model embeddings and PhyloP conservation scores.

| Model | HEK293FT | HMEC | HEPG2 | GM12878 | K562 | SKNSH |
|---|---|---|---|---|---|---|
| **DNABERT** | 0.22±0.01 | 0.44±0.01 | 0.35±0.01 | 0.35±0.01 | 0.28±0.01 | 0.26±0.01 |
| **DNABERT-3UTR** | 0.35±0.01 | 0.56±0.01 | 0.46±0.01 | 0.47±0.01 | 0.41±0.01 | 0.39±0.01 |
| **DNABERT-2** | 0.16±0.02 | 0.37±0.01 | 0.26±0.01 | 0.27±0.01 | 0.23±0.01 | 0.21±0.01 |
| **DNABERT2-ZOO** | 0.13±0.02 | 0.35±0.01 | 0.23±0.01 | 0.24±0.01 | 0.20±0.01 | 0.18±0.02 |
| **DNABERT2-3UTR** | 0.19±0.01 | 0.46±0.01 | 0.37±0.01 | 0.39±0.01 | 0.26±0.01 | 0.26±0.01 |
| **NT-MS-v2-250M** | 0.19±0.01 | 0.36±0.01 | 0.25±0.01 | 0.24±0.01 | 0.23±0.01 | 0.21±0.01 |
| **NTv2-250M-3UTR** | 0.31±0.01 | 0.51±0.01 | 0.42±0.01 | 0.41±0.01 | 0.34±0.01 | 0.32±0.01 |
| **StateSpace** | **0.40±0.01** | 0.55±0.01 | 0.50±0.01 | 0.51±0.01 | 0.40±0.01 | 0.39±0.01 |
| **StateSpace-SA** | **0.39±0.01** | **0.59±0.01** | **0.54±0.01** | **0.53±0.01** | **0.43±0.01** | **0.42±0.01** |
| **Griesemer et al., 2021** | 0.33±0.01 | 0.53±0.01 | 0.42±0.01 | 0.44±0.01 | 0.35±0.01 | 0.34±0.01 |

Table S4. Pearson r correlation coefficient between SVR-based predictions from sequence embeddings and ground truth MPRA activity from (Griesemer *et al.*, 2021).

| Model | Jurkat | | Beas2B | |
|---|---|---|---|---|
| | **steady state** | **stability** | **steady state** | **stability** |
| **DNABERT** | 0.20±0.02 | 0.31±0.02 | 0.21±0.02 | 0.37±0.03 |
| **DNABERT-3UTR** | **0.28±0.01** | **0.49±0.01** | **0.33±0.02** | **0.52±0.02** |
| **DNABERT-2** | 0.18±0.02 | 0.26±0.02 | 0.12±0.02 | 0.28±0.03 |
| **DNABERT2-ZOO** | 0.18±0.02 | 0.26±0.02 | 0.15±0.02 | 0.28±0.03 |
| **DNABERT2-3UTR** | 0.20±0.02 | 0.25±0.02 | 0.26±0.02 | 0.31±0.03 |
| **NT-MS-v2-250M** | 0.17±0.02 | 0.24±0.02 | 0.26±0.02 | 0.29±0.03 |
| **NTv2-250M-3UTR** | 0.25±0.01 | 0.35±0.01 | **0.34±0.02** | 0.46±0.02 |
| **StateSpace** | 0.26±0.01 | 0.35±0.01 | **0.31±0.02** | 0.43±0.02 |
| **StateSpace-SA** | **0.27±0.01** | 0.35±0.01 | 0.29±0.02 | 0.44±0.02 |
| **5-mers Siegel et al., 2022** | 0.22±0.01 | 0.43±0.01 | 0.15±0.02 | 0.48±0.02 |

Table S5. Pearson r correlation coefficient between Ridge-based predictions from sequence embeddings and ground truth MPRA data from (Siegel *et al.*, 2022).

| Model | Jurkat | | Beas2B | |
|---|---|---|---|---|
| | **steady state** | **stability** | **steady state** | **stability** |
| **DNABERT** | 0.23±0.01 | 0.36±0.01 | 0.22±0.02 | 0.40±0.03 |
| **DNABERT-3UTR** | **0.31±0.01** | **0.51±0.01** | **0.34±0.02** | **0.56±0.02** |
| **DNABERT-2** | 0.19±0.02 | 0.29±0.02 | 0.17±0.02 | 0.30±0.03 |
| **DNABERT2-ZOO** | 0.18±0.02 | 0.28±0.02 | 0.17±0.02 | 0.29±0.03 |
| **DNABERT2-3UTR** | 0.23±0.01 | 0.30±0.02 | 0.27±0.02 | 0.38±0.03 |
| **NT-MS-v2-250M** | 0.21±0.02 | 0.26±0.02 | 0.27±0.02 | 0.31±0.03 |
| **NTv2-250M-3UTR** | 0.28±0.01 | 0.39±0.01 | **0.36±0.02** | 0.48±0.02 |
| **StateSpace** | **0.31±0.01** | 0.38±0.01 | **0.32±0.02** | 0.48±0.02 |
| **StateSpace-SA** | **0.31±0.01** | 0.36±0.01 | **0.32±0.02** | 0.46±0.02 |
| **5-mers Siegel et al., 2022** | 0.25±0.01 | 0.45±0.01 | 0.19±0.02 | 0.48±0.02 |

Table S6. Pearson r correlation coefficient between SVR-based predictions from sequence embeddings and ground truth MPRA data from (Siegel *et al.*, 2022).

| Model | Ridge | SVR |
|---|---|---|
| **DNABERT** | 0.20±0.02 | 0.22±0.02 |
| **DNABERT-3UTR** | 0.28±0.02 | 0.29±0.02 |
| **DNABERT-2** | 0.32±0.02 | 0.35±0.02 |
| **DNABERT2-ZOO** | 0.30±0.02 | 0.36±0.02 |
| **DNABERT2-3UTR** | 0.28±0.02 | 0.33±0.02 |
| **NT-MS-v2-250M** | 0.31±0.02 | 0.34±0.02 |
| **NTv2-250M-3UTR** | **0.37±0.02** | **0.40±0.01** |
| **StateSpace** | **0.36±0.02** | **0.40±0.01** |
| **StateSpace-SA** | **0.33±0.02** | **0.39±0.01** |
| **k-mer (Agarwal and Kelly, 2022)** | 0.19±0.02 | 0.34±0.02 |

Table S7. Pearson r correlation coefficient between mRNA half-life prediction and ground truth data from (Agarwal and Kelley, 2022), using different 3'UTR embeddings.
